# The neural representation of personally familiar and unfamiliar faces in the distributed system for face perception

**DOI:** 10.1101/138297

**Authors:** Matteo Visconti di Oleggio Castello, Yaroslav O. Halchenko, J. Swaroop Guntupalli, Jason D. Gors, M. Ida Gobbini

## Abstract

Personally familiar faces are processed more robustly and efficiently than unfamiliar faces. The human face processing system comprises a core system that analyzes the visual appearance of faces and an extended system for the retrieval of person-knowledge and other nonvisual information. We applied multivariate pattern analysis to fMRI data to investigate aspects of familiarity that are shared by all familiar identities and information that distinguishes specific face identities from each other. Both identity-independent familiarity information and face identity could be decoded in an overlapping set of areas in the core and extended systems. Representational similarity analysis revealed a clear distinction between the two systems and a subdivision of the core system into ventral, dorsal and anterior components. This study provides evidence that activity in the extended system carries information about both individual identities and personal familiarity, while clarifying and extending the organization of the core system for face perception.

## Introduction

A wide and distributed network of brain areas underlies face processing. The model by Haxby and colleagues^1–3^ posited a division between a core system involved in the processing the visual appearance of faces—comprising the Occipital Face Area (OFA), the Fusiform Face Area (FFA), and the posterior Superior Temporal Sulcus (pSTS)— and an extended system, comprising parietal, frontal, and subcortical areas, involved in inferring socially relevant information from faces, such as direction of attention, intentions, emotions, and retrieval of person knowledge^1–4^.

The definition of the core system has been extended to include areas in the anterior fusiform gyrus (the anterior temporal face area, ATFA^5,6^), the anterior superior temporal sulcus (aSTS-FA^7–9^), and the inferior frontal gyrus (IFG-FA^7,10–12^). For example, in a recent fMRI neural decoding study with visually familiar faces^11^, we showed that the representation of face identity is progressively disentangled from image-specific features along the ventral visual pathway. While early visual cortex and the OFA represented head view independently of the identity of the face, we recorded an intermediate level of representation in the FFA in which identity was emerging but was still entangled with head view. The human face processing pathway culminated in the right ATFA and IFG-FA where we recorded a view-invariant representation of face identity.

While both unfamiliar and familiar faces effectively activate the core system^7,8,11,13,14^, familiar faces activate the extended system more strongly than unfamiliar faces^2,13,15–17^. Personally familiar faces recruit Theory of Mind (ToM) areas such as the medial prefrontal cortex (MPFC) and the temporo-parietal junction (TPJ), because they are more strongly associated with person knowledge^2,16,18^; they activate the precuneus and the anterior temporal cortices, suggesting retrieval of long-term episodic memories; they modulate the activity in the amygdala and insula, suggesting an increased emotion processing^2,13,18^. Because the core and extended systems have been mostly studied separately, we lack a clear understanding of how personal familiarity, consolidated through repeated interactions, affects the representations in the core system, and how core and extended systems interact to create the known behavioral advantages for personally familiar faces.

The behavioral literature on face processing^19–25^ suggests that, despite the subjective impression of efficient or “expert” perception of natural faces^26^, only familiar faces are detected and recognized more robustly and efficiently, in stark contrast with the surprisingly inefficient identification of unfamiliar faces. Recognition of personally familiar faces is highly accurate even when images are severely degraded, while recognition of unfamiliar faces is markedly impaired by variation in head position or lighting, even with good image quality^24,25,27–29^. Detection of personally familiar faces is facilitated even in conditions of reduced attentional resources and without awareness^19^.

The representations of familiar and unfamiliar faces may differ in multiple ways. Familiar identities could have more robust, individually-specific representations, which are learned and consolidated over the course of personal interactions. Alternatively, familiar face representations could be enhanced with attributes that are similar across many personally familiar faces. For example, personally familiar faces (especially those used in the present and our previous experiments that are faces of close relatives of personal friends) are associated with person-knowledge and emotional attachment that lead to social interactions that are different from the interactions with strangers, and these attributes may be shared across many familiar—one may be more open and unguarded with family and personal friends^18^.

Here we applied multivariate pattern analyses (MVPA^30,31^, including MVP classification (MVPC) and representational similarity analysis (RSA^32^ with two goals in mind. First, we wanted to dissociate familiarity information from identity information in the core and extended systems. Second, we wanted to investigate the relationships among core and extended face processing areas by examining the similarities of their representational spaces using second-order representational geometry^32–34^.

We first derived independent neural measures of identification and familiarity. To prevent any effect of familiarity information in identity decoding, we performed identity classification separately for familiar and unfamiliar faces. To control for the effect of identity-specific visual information in familiarity decoding, we trained classifiers to distinguish familiar from unfamiliar faces, and tested them on left-out identities. The results replicated the distinction between the representations of personally familiar and unfamiliar faces in the extended system that was previously revealed only with univariate analysis^2^, showing that this effect captured factors that were common across familiar faces and invariant across identities.

To unravel the representational structure of the face processing network, we investigated the relationships among the areas of the core and extended system uncovered by the classification analyses. Using the approach used by Guntupalli and colleagues^34^ (see also Kriegeskorte and colleagues^33^), we studied the similarities between representational geometries^32^ in different face-processing areas (second-order representational geometry). This analysis revealed clear distinctions between the core system and the extended system, supporting the model by Haxby and colleagues^1–3^. In addition, the results support the extension of the core system to more anterior areas, such as the ATFA, the aSTS-FA and IFG-FA^5–7,11,35^, and reveal a finer subdivision of this system into ventral, dorsal, and anterior components.

## Results

In this experiment, we investigated the face processing network while participants performed an oddball-detection task with faces of friends and strangers (see Figure 1). We first investigated which areas responded more strongly to familiar faces than unfamiliar ones with a standard GLM analysis. Because familiarity information (whether a face is a familiar one) is necessarily confounded with identity information (who that person is), we next used MVPC to dissociate which areas of the core and extended system encode identity-independent familiarity information (familiar vs. unfamiliar classification across identities), and which parts of the network encode identity information. We performed two classification analyses using different cross-validation schemes to control for the effect of identity on the representation of general familiarity and to control for the effect of familiarity on the representation of identity. For the familiarity classification, we employed a leave-two-identities-out cross-validation scheme, where the classifier was trained on six faces (three familiar, three unfamiliar) to distinguish between familiar and unfamiliar faces, and tested on two left-out identities. This cross-validation scheme reduced the effect of identity information (see Figure 4B and Supplementary Figures 1 and 2). For the identity classification, we decoded the four familiar faces and the four unfamiliar faces separately to eliminate the effect of familiarity information in the classification of identity information. Finally, we investigated the network structure derived from the similarities of representations to investigate relationships among areas in the core and extended system.

**Figure 1:**
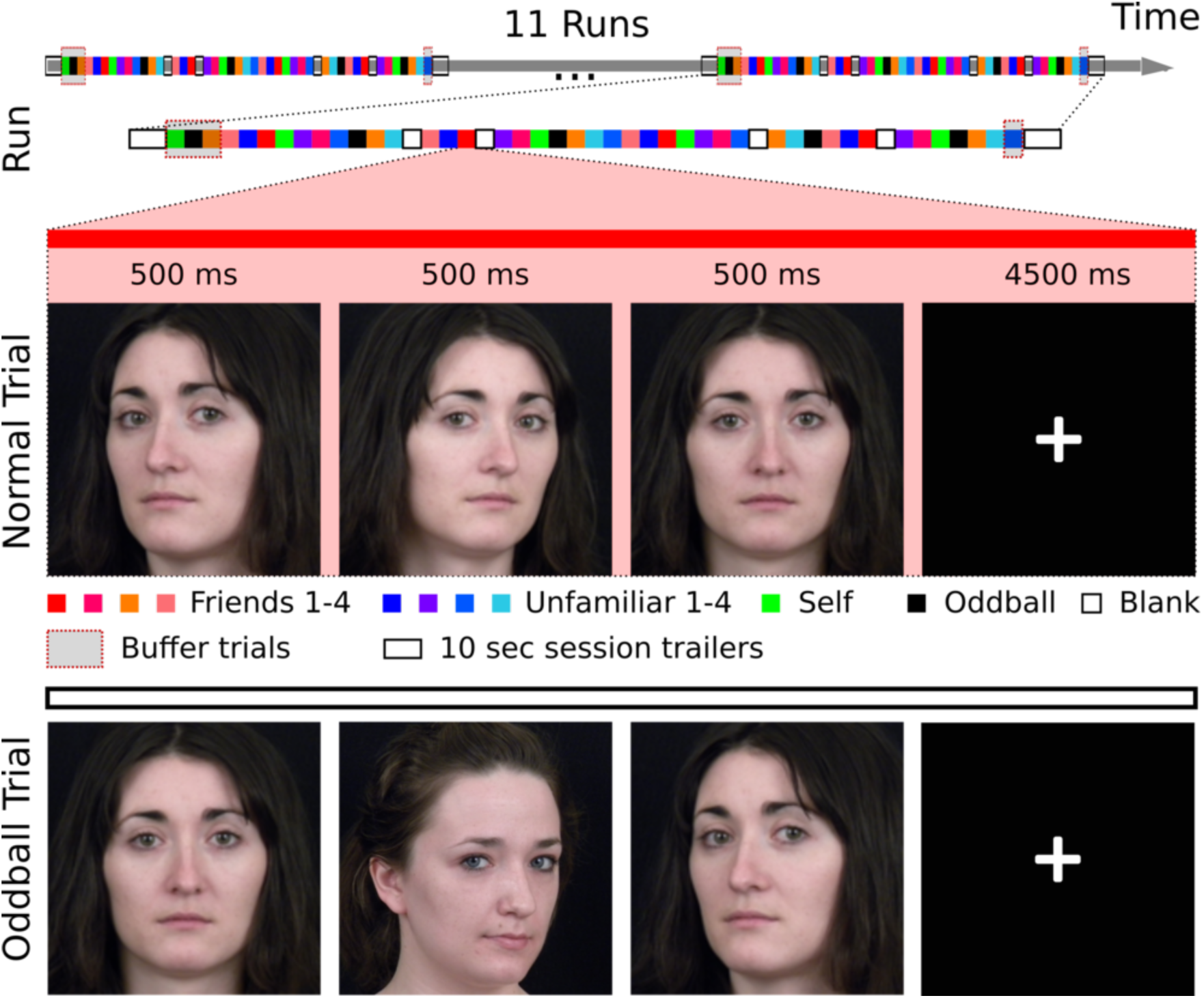
Slow event-related fMRI design. During each trial, images were presented in sequences of three pictures of the same identity (normal trial) or two different identities (oddball trials) in front-view or 30-degree profile views. Subjects engaged in an oddball-detection task to ensure that they paid attention to each stimulus.

**Figure 2.**
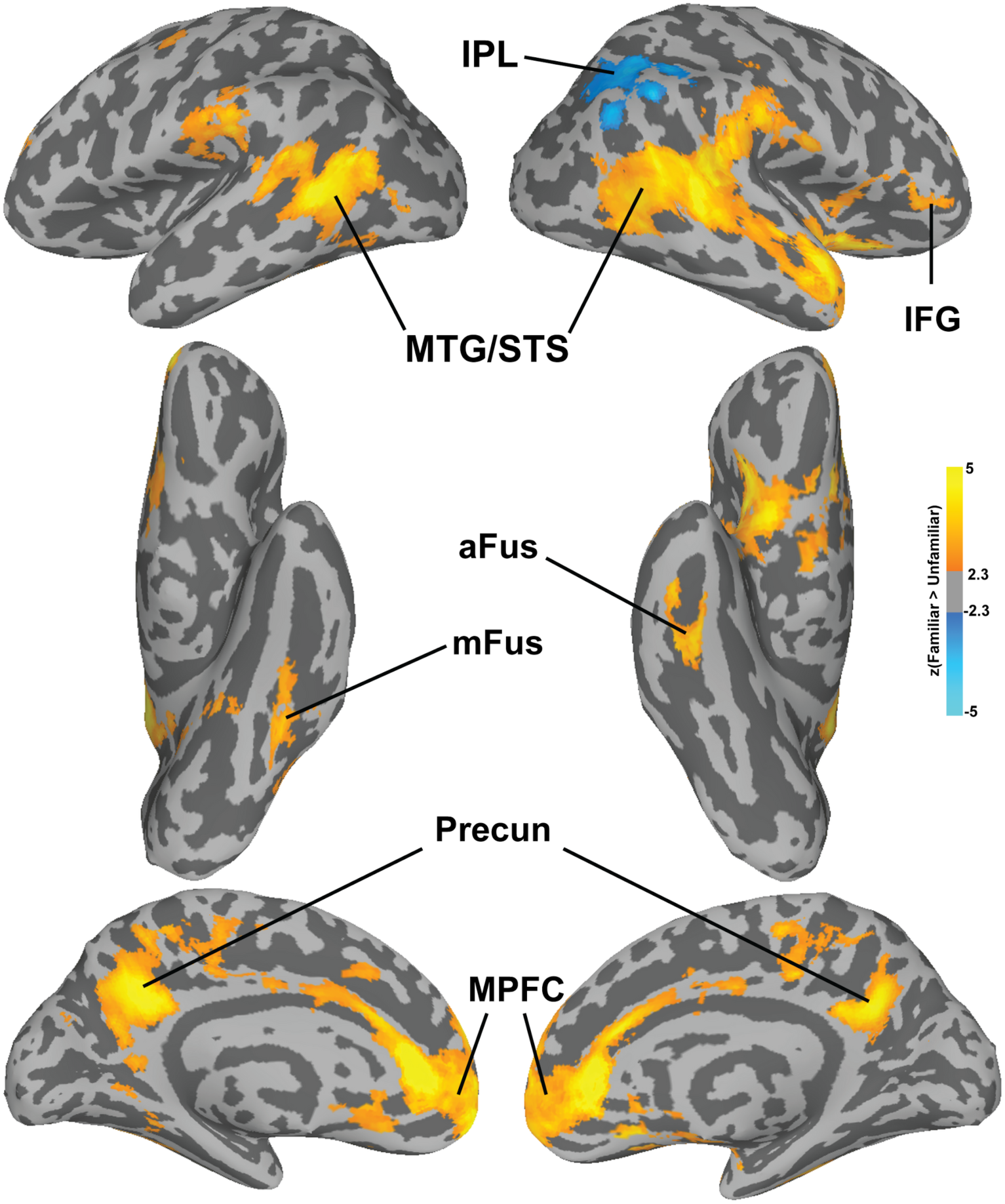
Cluster-corrected (p < .05) z-values for the univariate contrast Familiar > Unfamiliar. Abbreviations: IPL: inferior parietal lobule; mFus: middle fusiform gyrus; aFus: anterior fusiform gyrus; TPJ: temporo-parietal junction; MTG/STS: middle temporal gyrus/superior temporal sulcus; Precun: precuneus; MPFC: medial prefrontal cortex; IFG: inferior frontal gyrus.

### Univariate analyses

In the univariate analysis contrasting Familiar > Unfamiliar we found significant activation in bilateral MTG/STS extending along the full length of the right STS. Additionally, we found significant clusters in the bilateral precuneus and bilateral MPFC, as well as in the right IFG. Familiar faces also evoked stronger responses in the left mid fusiform gyrus and the right anterior fusiform gyrus near the locations of the FFA^36,37^ and ATFA^5^. For the contrast Unfamiliar > Familiar we found only one significant cluster in the right inferior parietal lobule encroaching on the TPJ. Figure 2 shows the resulting statistical maps projected on the surface.

## Multivariate analyses

### Familiarity Classification

The results of searchlight MVPC of identity-independent familiarity largely overlapped with the univariate maps, showing significant classification in the bilateral MTG/STS, mid and anterior right fusiform gyrus, right IFG, TPJ, precuneus, and MPFC (Figure 3). Surprisingly, small patches of cortex in early visual cortex also showed significant MVPC of identity-independent familiarity. We further investigated MVPC in early visual cortex with additional analyses on probabilistic ROI masks^38^, and found statistically significant decoding performance in V2 and V3 (see Supplementary Methods and Supplementary Figure 7). Since testing was performed on left-out familiar and unfamiliar identities, and all pictures were taken with the same equipment and settings, it is unlikely that this result was due simply to low-level features that distinguished familiar from unfamiliar faces. To test this further, we extracted features from the layers C1 and C2 of the HMAX model^39,40^ and performed the same classification analysis, and found that decoding performance was not statistically significant (accuracy with C1 features 52%, p = 0.67; accuracy with C2 features 49%, p = 0.96; see Supplementary Methods and Supplementary Figure 8).

**Figure 3.**
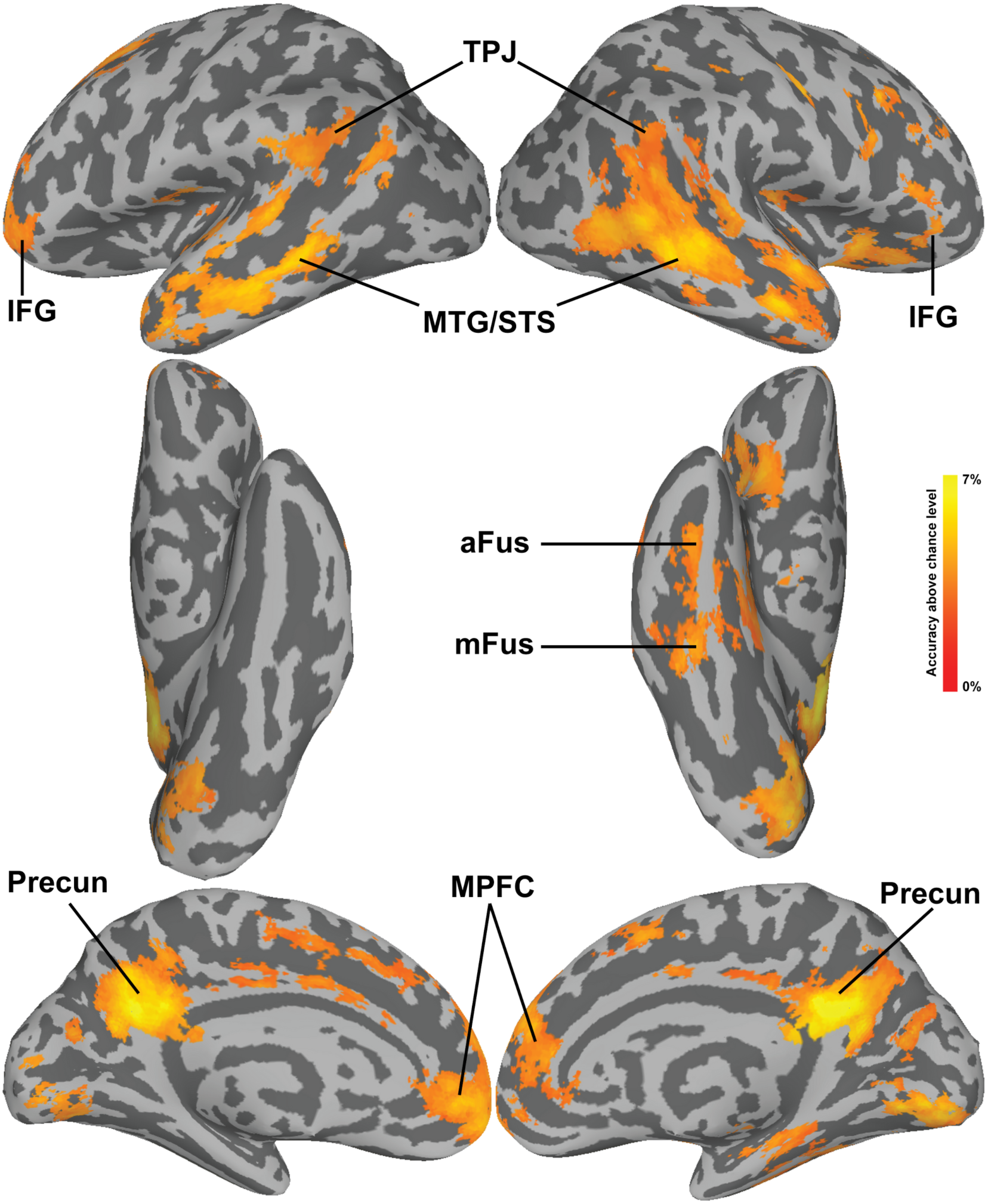
Searchlight maps for the Familiarity classification projected onto the surface. Maps were thresholded at a z-TFCE score of 1.65, corresponding to p < 0.05 one-tailed (corrected for multiple comparisons). Abbreviations: mFus: middle fusiform gyrus; aFus: anterior fusiform gyrus; TPJ: temporo-parietal junction; MTG/STS: middle temporal gyrus/superior temporal sulcus; Precun: precuneus; MPFC: medial prefrontal cortex; IFG: inferior frontal gyrus.

The results from the familiarity classification showed some correspondence with the univariate analysis, but the two maps were not completely overlapping. The scatterplot between the voxel-wise z-values for univariate GLM and the familiarity decoding MVPA results (Figure 4A) shows large spread, highlighting that the multivariate classification leveraged additional information other than magnitude differences between the two conditions. In addition, as can be seen in Figure 4B, the leave-two-identities-out cross-validation scheme effectively controlled for visual identity information in early visual cortex, which was necessarily conflated in the univariate contrast.

**Figure 4.**
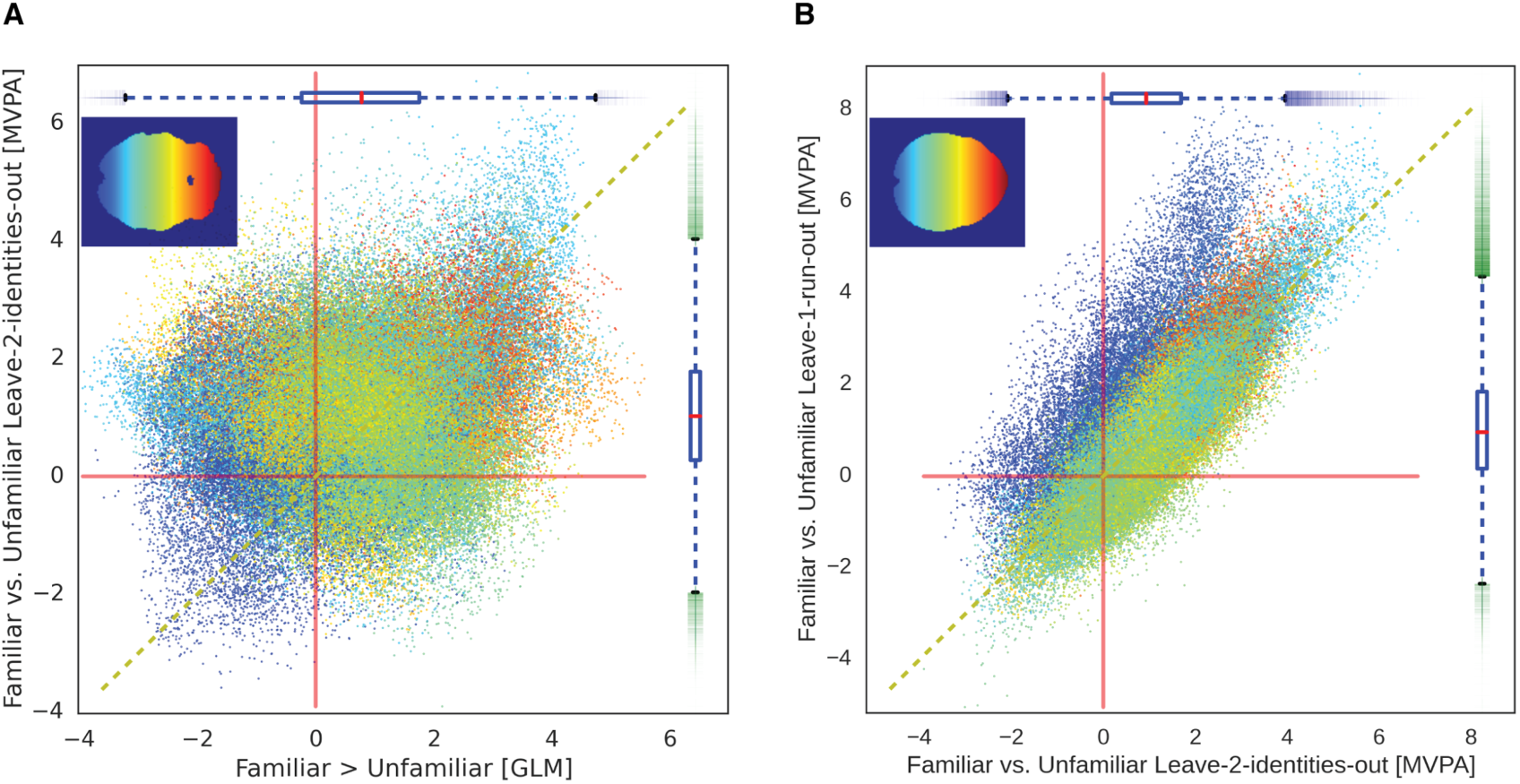
A) Comparison of the univariate analysis of familiarity with the MVPA familiarity decoding. The x-axis shows z-values from the univariate contrast Familiar > Unfamiliar. The y-axis shows the z-values of the Familiarity classification across identities. Colors depict voxel position in Posterior-to-Anterior (Blue-to-Red) direction. Some but not complete correspondence exists between the two maps, showing that the multivariate analyses leveraged additional information other than magnitude differences. **B) Comparison of cross-validation schemes for the familiarity decoding.** The x-axis shows z-values from the familiarity classification using the leave-2-identities-out scheme, as reported in the main manuscript. The y-axis shows the same classification using a common leave-one-run-out scheme. The leave-two-identities-out scheme successfully controls for identity visual information, as can be seen by the overall lower z-values for voxels belonging to the occipital cortex.

### Identity Classification

The identity classification analysis showed that identity could be decoded in many of the same areas as identity-independent familiarity (Figure 4). Significant classification was found in the MPFC and precuneus, and in the bilateral MTG/STS, TPJ, and IFG. The area in the precuneus with significant identity classification, however, was quite dorsal, whereas that for significant familiarity classification was ventral and included the posterior cingulate. Identity classification was significant in bilateral visual cortex starting in EV and extending to occipital, posterior, and mid fusiform cortices. Although MVPC of familiar identities showed a weak trend towards higher accuracies than for unfamiliar identities in the IFG and MTG/STS (Supplementary Figures 4, 5, and 6), these differences were not significant despite the large number of subjects.

**Figure 5.**
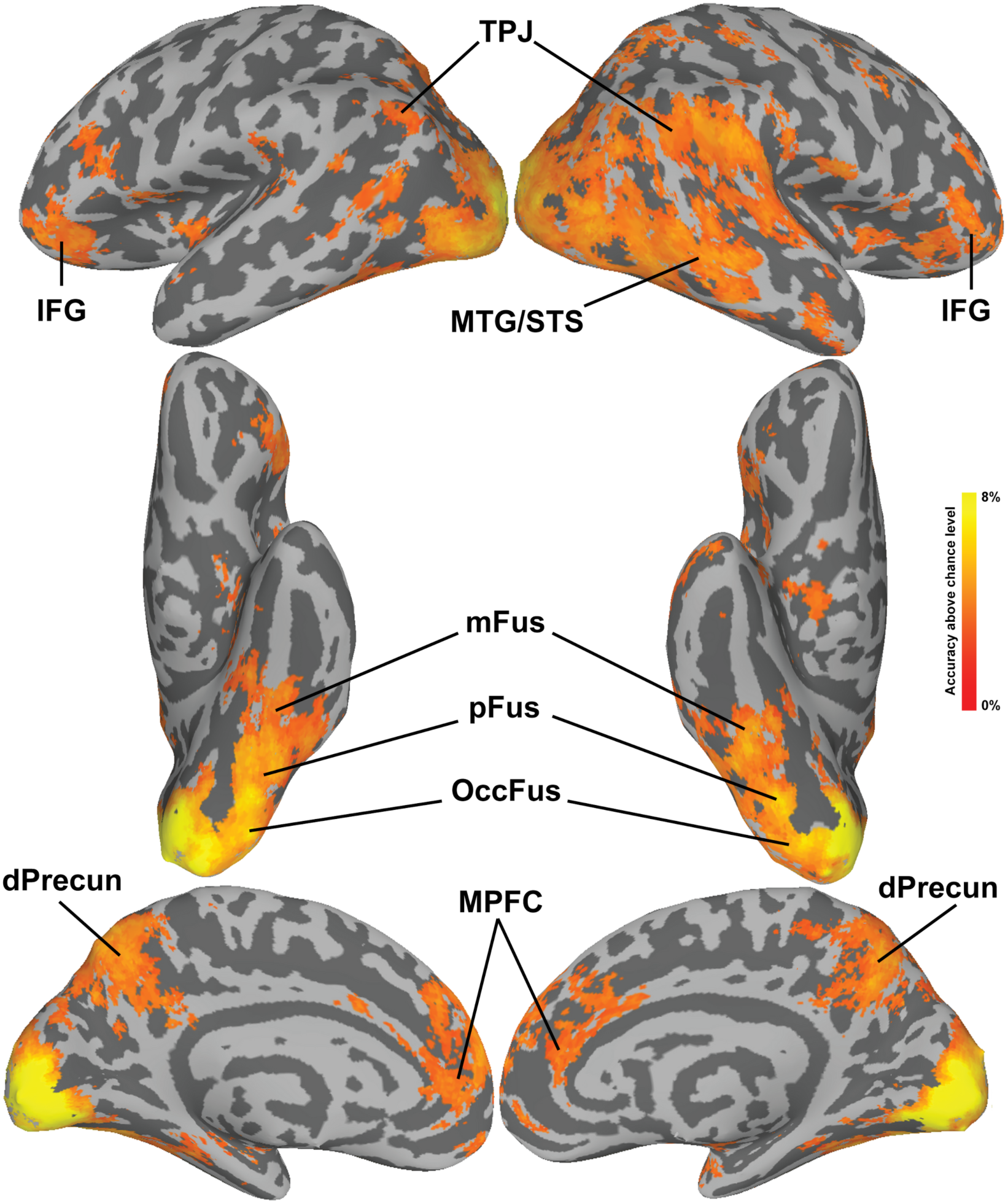
Searchlight maps for the Identity classification. The classification was run separately for familiar and unfamiliar identities (4-way), and the resulting maps were averaged. Maps were thresholded at a z-TFCE score of 1.65, corresponding to p < 0.05 one-tailed (corrected for multiple comparisons). Abbreviations: OccFus: occipital fusiform gyrus; pFus: posterior fusiform gyrus; mFus: middle fusiform gyrus; TPJ: temporo-parietal Junction; MTG/STS: middle temporal gyrus/superior temporal sulcus; dPrecun: dorsal precuneus; MPFC: medial prefrontal cortex; IFG: inferior frontal gyrus.

**Figure 6.**
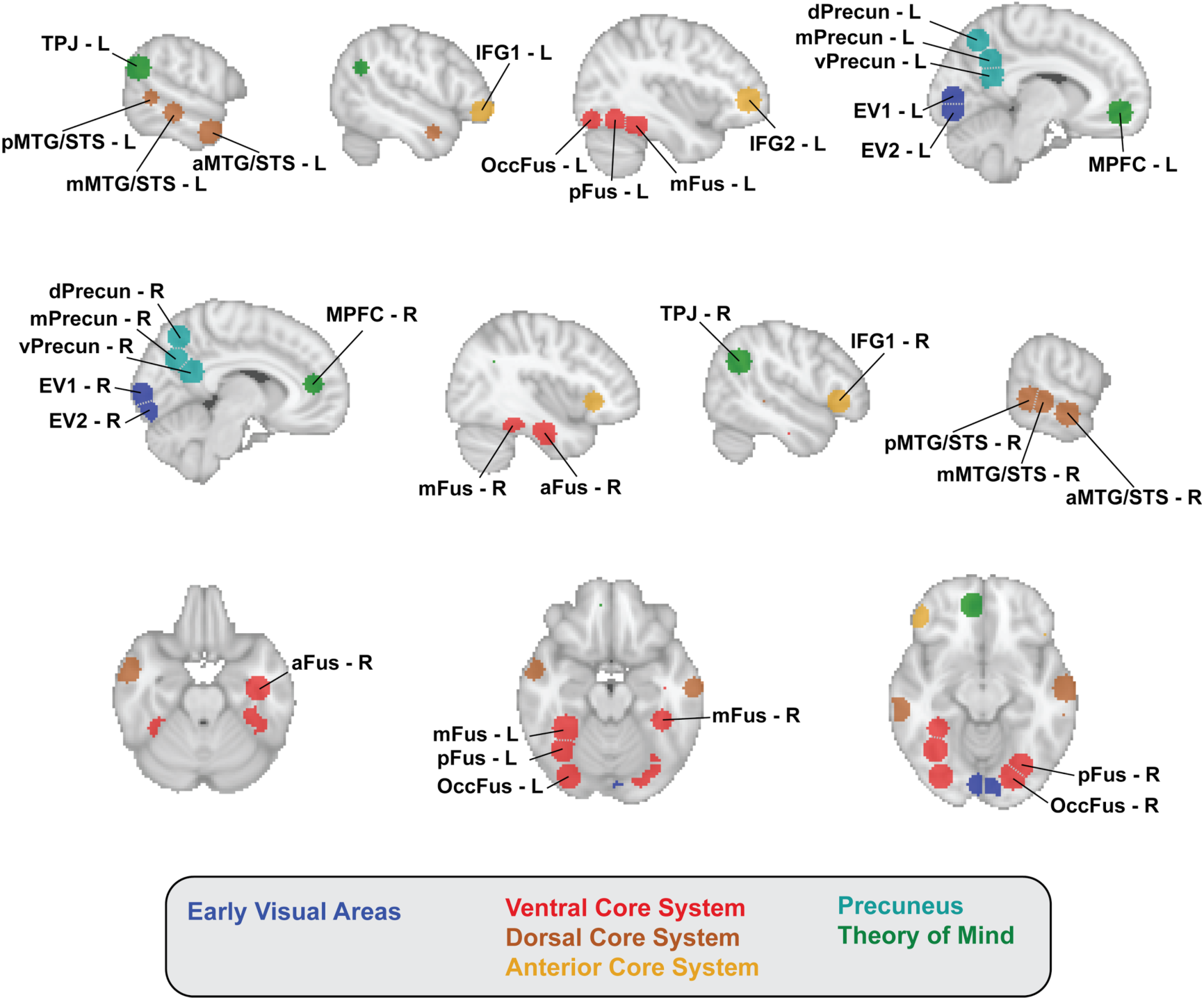
Spherical ROIs used to analyze the similarity of representational geometries. Top row shows left sagittal slices; middle row shows right sagittal slices; bottom row shows axial slices. Regions are color coded according to the system they belong to. Grey dotted lines between ROIs indicates that they were contiguous but not overlapping (see Methods for details).

### ROI Analysis and Second-order Representational Geometry

We investigated the relationships among the areas uncovered by the classification analysis as a second-order, inter-areal representational geometry. We selected 30 spherical ROIs (see Methods for how they were selected, Figure 5 for their location, and Supplementary Table 1 for their MNI coordinates) and computed a cross-validated representational dissimilarity matrix^41^ in each ROI. We then constructed a distance matrix quantifying the similarity of these RDMs between all pairs of ROIs. Then, we computed an MDS solution to visualize the geometry of this inter-ROI matrix. Figure 6 shows the results of the projection of a 3D MDS plot on the first two dimensions. Supplementary Figure 10 shows the distance matrices, Supplementary Figures 11, 12, and Supplementary Videos 1, 2 show the full MDS solution.

The first two dimensions of the MDS solution captured relationships among areas in the ventral portion of the core system in the first dimension, and relationships among areas in the dorsal and anterior parts of the core system and areas in the extended system in the second dimension. The first dimension showed a progression from EV areas to the posterior, mid, and anterior fusiform areas. Extended system areas were all at the distant end of the first dimension, as were the areas in the dorsal part of the core system (MTG/STS) and the IFG. The second dimension captured distinctions among these extended and core system areas, with the precuneus areas clustered together at one end, the MPFC and TPJ in the middle, and the dorsal and anterior core system areas at the other end.

We replicated this second-order RSA on an independent fMRI dataset collected while different subjects watched a full-length audiovisual movie, *Raiders of the Lost Ark*^*34,42*^ This naturalistic stimulus contained a rich variety of dynamic faces that rapidly became familiar while the plot unfolded. The inter-ROI similarity matrix and MDS plot replicated the results based on representational geometry for the eight faces in the experiment (Figure 6). The results tend to be more clearly defined for the movie data, probably due to the the dynamic videos, the larger data set, and hyperalignment of the data. Contributions from scene context, language, music, and narrative structure might also play a role^43,44^. The first two dimensions of the MDS solution cleanly captured distinctions in the ventral core system in the first dimension and in the extended, dorsal core, and anterior core systems in the second dimension, with remarkably similar placement of ROIs on each of these dimensions between task data and movie data; the distance matrices obtained from the two datasets were very similar (RV-coefficient^45,46^ = 0.755 [0.7254, 0.7612]; Spearman r = 0.48 [0.34, 0.49]; see Supplementary Methods and Figures 12, 13).

We quantified the similarity of the within-system RDMs by running a linear mixed-effect model on the correlation values and contrasting within-systems correlations with between-systems correlations. We found a clear distinction between the core and extended systems in terms of similarity of representational geometries. For the task data, the correlations within the extended system were significantly higher than the between-system correlations (estimate of the contrast “Within Extended > Between” 0.0993 [0.0875, 0.1111] 95% confidence interval, t-value = 16.36), while the correlations within the core system were not significantly different from the between-system correlations (estimate of the contrast “Within Core > Between” 0.0044 [- 0.0043, 0.0130], t-value = 1.00). For the movie data, both contrasts were significant: within-core vs. between 0.0678 [0.0619, 0.0738], t-value = 22.47; and within-extended vs. between 0.1479 [0.1398, 0.1565], t-value = 35.07. Supplementary Tables 2 and 3 show the full parameter estimates for both models, while Supplementary Tables 4 and 5 report additional statistics on the subsystems.

**Figure 7.**
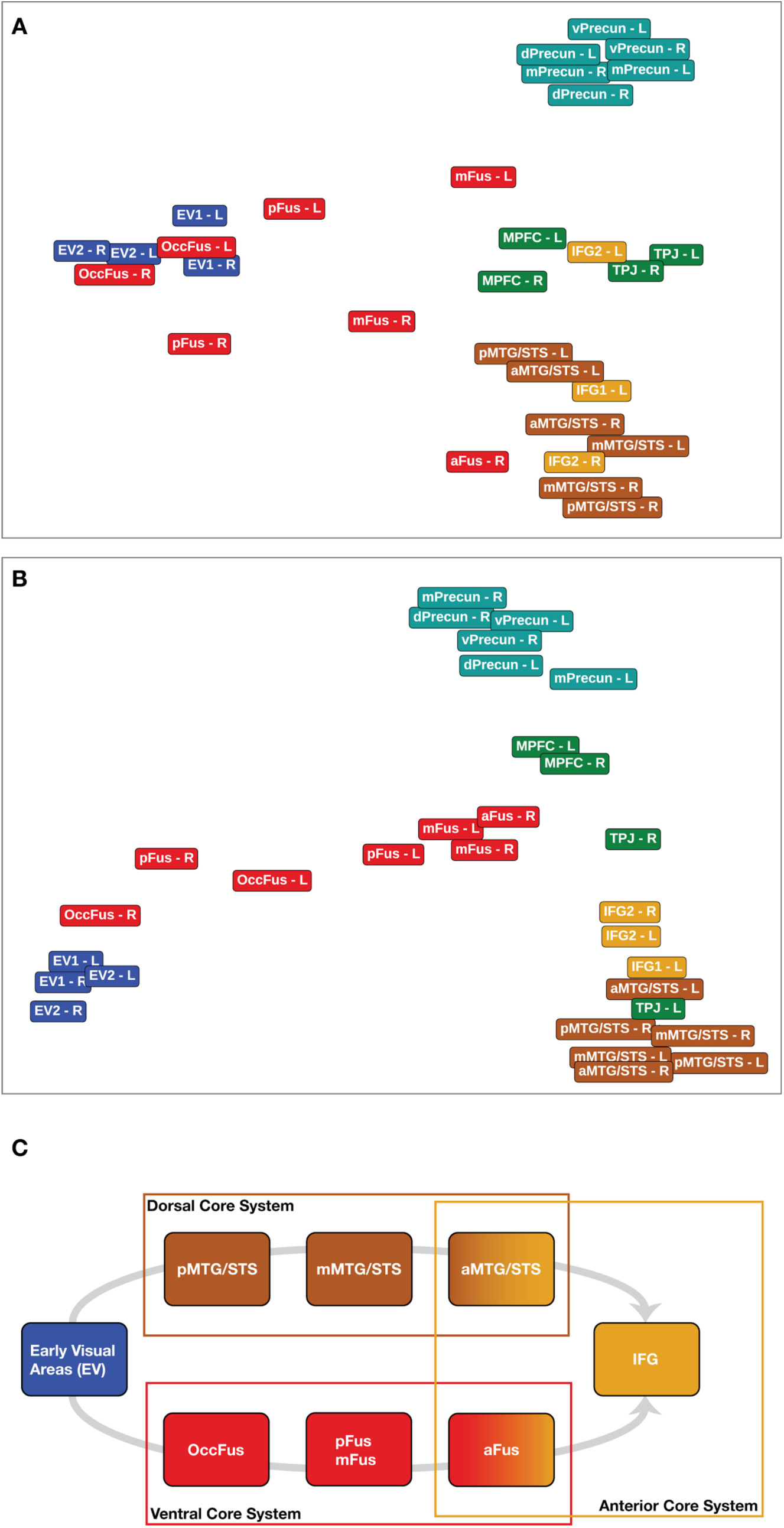
Similarity of neural representations in ROIs derived from familiarity and identity decoding. Top panel and middle panel show the first two dimensions of 3D MDS solutions based on the task data (A) and the hyperaligned movie data^34,42^ (B) (see Methods section for more details). The color of the labels indicates the system to which the ROI belongs to (see Figure 5 for their location and Supplementary Table 1 for the MNI coordinates). With both datasets the MDS solution shows the hierarchy from early visual cortex to ventral core system (first dimension, x-axis), as well as a segregation between the precuneus, theory of mind areas, and areas of the anterior and dorsal core system (second dimension, y-axis). Panel (C) shows the proposed division of the core system into dorsal, ventral, and anterior portions. This division builds upon existing models of face processing^1,7^, which suggest a progression from early visual areas to separate dorsal and ventral streams, while extending these models with the finding that both streams converge to the IFG. Representation of identity and gaze in the anterior core areas are disentangled from variations in head view^11,47^.

## Discussion

In this experiment we investigated how familiar and unfamiliar faces are represented in the distributed neural system for face perception. We distinguished between familiarity information, abstracted from the visual appearance of the faces, and the identification of individual faces, controlling for the added information of personal familiarity. These analyses revealed an extensive network of areas that carry information about face familiarity and identity, replicating with a larger sample size previous studies that used univariate analyses, and also providing more details about the type of information present in those areas. We then analyzed the second-order representational geometry of this extensive network, revealing a clear distinction between the core and the extended systems for face perception and a new subdivision of the areas in the core system.

The results from the second-order representational analysis suggest that the core system for face perception can be separated into ventral, dorsal, and anterior subsystems, extending the existing neural models of face perception^1,2,7^. The ventral core system consists of fusiform areas extending from the occipital lobe to the anterior ventral temporal lobe. The dorsal system extends from the posterior MTG/STS to anterior lateral temporal cortex. The representations in the dorsal core system did not appear to have strong similarities with those in the ventral core system, consistent with the functional distinction between dorsal and ventral areas suggested by others^48, 8^. Based on the results reported here, we propose that the anterior areas in the fusiform gyrus, the anterior MTG/STS, and the IFG may be the convergence of the ventral and dorsal pathways in which representations of faces become invariant to facial attributes such as head position^9,11^ and perhaps other social attributes. For example, the right anterior STS plays a role in the representation of the dangerousness of animals^49^ and may play a role in the representation of social impressions, such as trustworthiness and aggressiveness^50^. The hierarchy of areas proposed in this work provides a testable model for future studies aimed at further characterizing the transformations operating on the representation of faces, from retinotopic input to higher order areas.

Using MVPC, we teased apart neural responses due to factors that are shared by familiar faces from factors that are specific to familiar and unfamiliar identities. While standard univariate analyses necessarily conflate identity information with familiarity information, we used different cross-validation schemes in the MVPC to separate familiarity information from identity information. To separate identity-independent familiarity information from identity-specific visual information, we used a cross-validation scheme in MVPC of face familiarity in which we tested the classifier on identities that were not included in the training data. To investigate identity-specific information that was independent of familiarity, we tested MVPC of familiar and unfamiliar identities separately.

We found reliable decoding of identity-independent familiarity in extended system areas that showed stronger responses to familiar faces in univariate analyses, such as theory of mind areas (precuneus, TPJ, and MPFC), consistent with previous reports^2,13^. Univariate and multivariate analyses were complementary in that they tested different properties of the BOLD response (differences in mean activations vs. patterns): the two resulting maps showed some but not complete correspondence, highlighting that multivariate analyses leveraged additional information other than magnitude differences. In addition, MVPC of familiarity was designed to test for a familiarity effect that was not specific to familiar individuals, revealing that this network does carry such identity-independent information about the familiarity of faces. Both the univariate and MVPC results expand the areas reported previously to include additional areas that are components of the dorsal and anterior core system for face perception in the MTG/STS, anterior fusiform cortex, and IFG. We suspect that our relatively large sample size made it possible to identify this more extensive network.

In this experiment subjects had to perform an oddball-detection task to ensure that they paid attention to the stimuli. It is possible that some of the decoding results for familiarity might be attributed to differences in attentional demands between personally familiar and unfamiliar faces, but it is hard to predict the direction of an effect of attention. Behavioral evidence suggests that personally familiar faces are processed faster^20–23,51^, and require fewer attentional resources^19^. On the other hand, we also have shown that familiar faces, relative to unfamiliar faces, slow down shifts of attention away from the face, suggesting they hold attention^52^. We found reduced BOLD activation to personally familiar faces only in the IPL^53,54^, while areas of the core and extended systems showed stronger responses. If the stronger response to familiar faces in core and extended system areas were due to spontaneous attention, one would also expect a stronger response in the IPL and other attention-related cortical areas, which we did not find.

Unexpectedly, we found significant decoding of familiarity information in early visual cortex while controlling for identity information. Additional ROI decoding analyses in early visual areas^38^ revealed that familiarity information could be decoded in V2 and V3 (see Supplementary Material). Low-level image differences did not seem to explain this finding: familiar and unfamiliar faces were indistinguishable using features extracted from the HMAX model^39,40^. Recent studies have shown that feedback information from higher-order visual areas to early visual cortex carries fine-grained information about the category of the stimuli being observed^55,56^, suggesting that feedback processes might have contributed to the significant familiarity decoding in early visual areas. However, future studies with paradigms designed to address the nature of these feedback processes are needed to further test this possibility.

In addition to identity-independent familiarity, the same network carries information about specific identities. We tested for this type of information with separate MVPC analyses of four familiar identities and four unfamiliar identities. By not including familiar and unfamiliar identities in the same analysis, we could test for identity-specific neural patterns that were not dependent on familiarity. Again, this network was more extensive than that reported in previous studies^11,57–63^ (most probably due to the larger number of subjects and, perhaps, the inclusion of personally familiar faces). Importantly, this network included the IFG, consistent with previous work^11,12^, and extended into the MTG/STS, TPJ, precuneus, and MPFC.

Identity decoding was also found in early visual cortex and the posterior ventral core system, likely reflecting to some extent image-specific information. In a recent experiment^11^ we showed that view-dependent representation of faces was the dominant factor in early visual cortex and the OFA. We did not find a significant difference in MVPC of familiar identities as compared to MVPC of unfamiliar identities, despite the large number of subjects in this study. There was a nonsignificant trend towards higher MVPC accuracies for familiar identities in the IFG and MTG/STS, but more work explicitly designed to investigate view-invariant representations of identity is needed to establish whether these trends are real.

## Conclusions

Our results revealed new structure in the distributed system for face perception, suggesting that the core system can be subdivided into ventral, dorsal, and anterior components based on differences of representations. The anterior portion of the core system may be the point at which the ventral and dorsal pathways converge to generate view-independent representations of identity and of socially-relevant visual information, such as direction of attention. Identity-independent information about familiarity could be decoded in extended system areas such as the TPJ, precuneus, and MPFC, as well as in dorsal and anterior core system areas such as the MTG/STS, anterior fusiform cortex, and IFG. In sum, these results reveal new information about how face perception, one of the most highly developed and socially relevant visual functions, is realized in an extensive distributed system involving cortical fields in occipital, temporal, parietal, and prefrontal cortices.

## Materials and Methods

### Participants

Thirty-three young adults participated in the experiment (mean age 23 y.o. +/- 3.33 SD, 13 males). They were recruited from the Dartmouth College community and all had normal or corrected-to-normal vision. Prior to the imaging study we took pictures of four friends for each participant to use as familiar stimuli. Some of these friends also were study participants (pictures of 76 individuals were taken as familiar stimuli). Photos of unfamiliar individuals were collected at the University of Vermont (Burlington) using the same camera and lighting conditions. All individuals signed written informed consent to use their pictures for research and in publications. Prior to participation in the fMRI study, subjects were screened for MRI compliance and provided informed consent. The study was approved by the Committee for the Protection of Human Subjects at Dartmouth College and was conducted according to the principles of the Declaration of Helsinki. Participants received monetary compensation for their time.

### Stimuli

The stimuli for the fMRI experiment were pictures portraying different familiar and unfamiliar identities: four friends’ faces, four unknown faces, and the subject’s own face. For each identity we used three images with different head orientations: frontal view and 30-degree profiles to the left and right with gaze towards the camera. All photos on both sites (Dartmouth College and University of Vermont) were taken using the same consumer-grade digital camera in a dedicated photo-studio room with black background and uniform lighting.

Each familiar face was matched with an unfamiliar individual face, similar in age, gender and ethnicity. Twenty-seven images (9 individuals, 3 head positions) were used in the experimental design per each subject. Stimuli were presented to the subjects in the MRI scanner using a projection screen positioned at the rear of the scanner and viewed through a mirror mounted on the head coil.

The original high-resolution digital images were cropped to include the face from the top of the head to the neck visible under the chin, centered on the face. Images were scaled to 400x400 pixels. Images subtended approximately 10x10 degrees of visual angle.

### Procedure

The stimuli were presented using a slow event-related design while subjects were engaged in a simple oddball task (Figure 1). A typical trial consisted of three different images of the same individual, each presented for 500 ms with no gap. On catch trials, one of the three images was of a different individual. The order of head orientations within trials was randomized. The task was included to make sure that subjects paid attention to the identity of the faces. Before entering the scanner, subjects had a short practice session with each condition (one trial for each of 9 identities, one blank trial, and one catch trial) to be familiarized with the design and the stimuli.

The order of the events was pseudo-randomized to approximate a first-order counterbalancing of conditions^64^. A functional run comprised 48 trials: four trials for each of the nine individuals (four familiar, four unfamiliar and self), four blank trials, four oddball and four buffer trials (three at the beginning and one at the end). The buffer trials were added to optimize the trial order and were discarded from the analysis. Each run had 10 seconds of fixation at the beginning (to stabilize the hemodynamic response) and at the end (to collect the response to the last trials). Each session consisted of 11 functional runs, resulting in 396 non-oddball trials (44 for each of the nine identities).

### Image acquisition

Brain images were acquired using a 3T Philips Achieva Intera scanner with a 32-channel head coil. Functional imaging used gradient-echo echo-planar-imaging with SENSE reduction factor of 2. The MR parameters were TE/TR = 35/2000 ms, Flip angle = 90°, in-plane resolution = 3×3 mm, matrix size of 80×80 and FOV = 240×240 mm. 35 axial slices were acquired with no gap covering the entire brain except the most dorsal portion (Supplementary Figure 9). Slices were acquired in the Philips-specific interleaved order (slice step of 6, i.e., ceiled square root of total number of slices). Each of the 11 functional runs included 154 dynamic scans with 4 dummy scans for a total time of 316 seconds per run. After the functional runs a single high-resolution T1-weighted (TE/TR = 3.7/8.2 ms) anatomical scan was acquired with a 3D-TFE sequence. The voxel resolution was 0.938×0.938×1.0 mm with a bounding box matrix of 256×256×160 (FOV = 240×240×160 mm).

### Image preprocessing

All preprocessing steps were run using a Nipype workflow (version 0.11.0; FSL version 5.0.9)^65,66^, which also used functions from SciPy^67^ and NumPy^68^. We modified the preprocessing pipeline *fmri_ants_openfmri.py* and adapted it for our analyses. The modified version is available at https://www.github.com/mvdoc/famface. All the preprocessing analyses were run on a computing cluster running Debian Jessie with tools provided by the NeuroDebian repository^69^.

### Preprocessing Steps

We used a standard FSL preprocessing pipeline (FEAT) as implemented in Nipype (*nipype.preprocess.create_featreg_preproc*), using a FWHM smoothing of 6 mm, a highpass filter at 60 s cutoff, and the first volume of the first run as a reference for EPI alignment. After motion correction, the BOLD time-series were masked with a dilated gray-matter mask, smoothed, and then high-pass filtered. The preprocessed data were then used for a GLM and MVPA analysis, with additional preprocessing steps as described in the following sections.

#### Template Registration

Each subject’s data (functional or second-level betas) were resliced into the MNI template with 2 mm isotropic voxel size. First, a reference volume was created by computing a median temporal SNR volume across functional runs. Then, we computed an affine transformation registering this median tSNR volume to the subject’s anatomical scan using FSL’s FLIRT tool^70^, and the transformation was improved using the BBR cost function. A second non-linear transformation registering the subject’s anatomical image to the MNI template was computed using ANTs^71^ with default parameters. The affine and nonlinear transformations were then combined to reslice the reference volume and all the functional volumes and second-level betas into the MNI template. Results from this registration pipeline were visually inspected for each subject.

#### MVPA Preprocessing

First, we resliced the bold time-series into the MNI template using a combination of linear and nonlinear transformations (see Template Registration section). Then, we extracted beta parameters associated with each condition for each run using PyMVPA’s *fit_event_hrf_model* ^72^ function based on NiPy’s functionality^73^. Additional nuisance regressors comprised motion estimates, artifacts (volumes were marked as artifact if their intensity exceeded three standard deviations of the normalized intensity), and noise estimates. To obtain noise estimates we used the CompCor method^74^. In brief, we performed a GLM on the BOLD timeseries in the voxels belonging to each subject’s white-matter mask projected in MNI space. The regressors of this GLM were the motion estimates and volumes marked as artifacts. We then performed PCA on the residuals, and took the first 5 components as noise estimates.

### Univariate analyses

The first-level and second-level analyses (fixed effect) for each subject were performed in the subject’s individual space, and the results were then projected into a standard template (FSL’s MNI152, 2 mm isotropic, see details in the Template Registration section). These analyses followed a standard FSL pipeline as implemented in Nipype (*nipype.estimate.create_modelfit_workflow* and *nipype.estimate.create_fixed_effects_flow*). A standard GLM analysis was performed separately for each run to extract beta values associated with each condition and the planned contrasts. Additional nuisance regressors comprised motion estimates, artifacts (volumes were marked as artifact if their intensity exceeded three standard deviations of the normalized intensity), and first-order derivatives. A second-level analysis was performed to obtain per-subject statistical maps associated with each condition and contrast using FSL’s *FLAMEO* (fixed-effect model). The statistical maps were then resliced into the MNI152 template (see details above), and a third-level analysis was performed across subjects using FSL’s *FLAMEO* (mixed-effect model). The resulting z-stat maps were then corrected for multiple comparisons using FSL’s *cluster* routine, with a voxel z-threshold set at 2.3, and cluster p-value of p = .05. The Nipype pipeline we used for third-level analysis can be found at https://www.github.com/mvdoc/famface^1^.

## Multivariate analyses

### Classification methods

MVPC was implemented in Python using PyMVPA^72^ http://www.pymvpa.org). GLM betas were estimated within each run for each condition (see MVPA Preprocessing section). For all analyses we kept only the betas for the four familiar and the four unfamiliar identities, discarding trials where subjects saw their own face, or responded to an oddball presentation. The betas were then z-scored within each run (separately for each voxel) and used as features for classification. We used Linear C-SVM as a classifier, as implemented in LIBSVM^75^. The C parameter was set to the PyMVPA default, which scales it according to the mean norm of the training data.

### Cross-validation

We used a leave-one-out (LOO) scheme for cross-validation. The splitting unit was dependent on the type of classification (familiarity or identity). For familiarity classification, we cross-validated across pairs of identities. We trained the classifier on three familiar and three unfamiliar identities, and tested on the left-out identities. This resulted in 16 cross-validation splits that allowed us to control for identity information (see Supplementary Figures 1 and 2 for a comparison of leave-one-run-out and leave-two-identities-out cross-validation schemes). For identity classification, we cross-validated across runs, resulting in a leave-one-run-out scheme (11 splits). To remove the effect of familiarity on classification of face identity, we performed identity classification independently for familiar and unfamiliar identities, and averaged the resulting accuracy maps.

### Searchlight

We used sphere searchlights^76^ to extract local features for classification. We selected a 5-voxel radius (10 mm), and moved the searchlight sphere across the voxels belonging to a union mask in which at least 26 subjects (∼80%, arbitrarily chosen) had fMRI coverage (see Supplementary Figure 9), as well as selecting only gray- and white-matter voxels in the cerebrum. For each center voxel in this mask, we selected nearby voxels contained in a sphere, and used them as features for classification. The classifier’s accuracy was stored in the central voxel, and the process was repeated for every voxel.

### Statistical assessment

To determine statistical significance for the MVPC analyses, we performed permutation testing^77^ coupled with Threshold-Free Cluster Enhancement (TFCE)^78^, as implemented in CoSMoMVPA^79^. For each subject and each classification analysis, we computed a null distribution by randomly permuting the labels and performing classification. For identity classification analysis, we randomly shuffled the identity labels within each run, and performed classification. This procedure was repeated 20 times for each subject. For familiarity analysis, we randomly permuted the familiarity labels across the entire experiment. This was repeated exhaustively, resulting in 35 permutations (see Supplementary Materials for a short proof that only 35 unique permutations are possible in this case). To create a null distribution of TFCE values for each voxel, permutation maps were randomly sampled and averaged across subjects, and this process was repeated 10,000 times. Note that we selected a smaller number of permutations than suggested^77^ (100 per subject) because of the large number of subjects we had: with 33 subjects, the number of possible average maps for identity classification was 20^33^ and for familiarity classification was 35^33^.

### Similarity of neural representations within ROIs

#### Second-order Representational Similarity Analysis

We defined ROIs based on the searchlight results for both the familiarity and identity classification. Thirty spherical ROIs were centered on voxels selected manually at or near peak values, with a 10 mm radius (five voxels). Voxels belonging to more than one ROI were assigned to the ROI with the closest center (Euclidean distance), resulting in some contiguous but not overlapping ROIs (see Figure 5). On average, ROIs contained 412 voxels at a 2 mm isotropic resolution (SD: 73 voxels).

For each ROI we computed a cross-validated representational dissimilarity matrix (RDM)^41^ between the eight identities (four familiar faces, four unfamiliar faces). First, we z-scored the beta estimates within each run, which were computed as described in the MVPA Preprocessing section. Then, we divided all runs into two partitions of six and five runs, and averaged the beta values within each partition. The data between these two partitions were correlated (Pearson correlation) to obtain an 8x8 matrix of dissimilarities between pairs of identities. Note that because correlations were computed between data from two different partitions, the diagonal could be different from one. This process was repeated for every possible combination of runs, yielding 462 RDMs that were averaged to obtain a final RDM for each ROI and each subject. The final RDMs were made symmetrical by averaging them with their transpose. All averaging operations were performed on Fisher-transformed (r-to-z) correlation values, then mapped back to correlation using the inverse transformation.

We used these final RDMs to compute pairwise distances between ROIs for each subject individually using correlation distance. The resulting 33 distance matrices (one for each subject) were averaged to obtain a group-level distance matrix. This distance matrix was used to compute a three-dimensional MDS solution, using classical MDS as implemented in R (*cmdscale*) interfaced in Python using *rpy2*^*80*^.

#### Comparison with movie data

To investigate the reproducibility of the network formed by the ROIs defined above, we computed between-subject correlation distances across these ROIs using hyperaligned data from a different study, in which eleven participants watched “Raiders of the Lost Ark”^34,42^. Since data were functionally aligned with hyperalignment^34,42^, we performed a between-subject analysis instead of a within-subject analysis, where distances between pairwise ROIs were computed across subjects, replicating the approach in Guntupalli et al.^34^. Additional details on the experimental paradigm and scanning parameters can be found in the Supplementary Material.

Because data were in two different resolutions of the same template (task: MNI 2 mm; movie: MNI 3 mm), center coordinates of the spherical ROIs were recalculated assigning the closest voxel in MNI 3 mm using Euclidean distance. The median displacement was 1.41 mm (min: 1 mm, max: 1.73 mm). As described above, spherical ROIs were drawn around these center voxels using a radius of 9 mm (3 voxels) to account for the different voxel size. Overlapping voxels were assigned to the ROI with the closest center, resulting in possibly contiguous but not overlapping ROIs. On average ROIs contained 100 voxels (SD: 20 voxels).

The movie data were masked selecting only white- and gray-matter voxels, and divided into two parts for cross-validation. For each of the two parts, whole-brain searchlight hyperalignment parameters were derived from one part of the movie, and the second part was projected into the common model space in functional alignment ^34,42^. The aligned data were z-scored, and timepoint-by-timepoint RDMs were computed in each ROI for each subject individually, yielding a 1322 x 1322 RDM within each ROI (1336 x 1336 for the second fold of hyperalignment). Following the analysis in (Guntupalli et al., 2016) we estimated a distance matrix between ROIs while cross-validating across subjects. For each pair of ROIs, the correlation between their RDMs was computed for all 55 pairs of subjects, and averaged to compute the cross-validated correlation between those ROIs. This process resulted in two 30x30 cross-validated distance matrices (one for each hyperalignment fold), which were made symmetrical by averaging them with their transpose, and finally averaged together to obtain one final 30x30 matrix. All averaging operations were computed on Fisher-transformed (r-to-z) correlation values, then mapped back to correlation using the inverse transformation. Finally, a dissimilarity index (D) was computed for each pair of ROIs to normalize the correlation according to the maximum possible correlation within each ROI^33^:

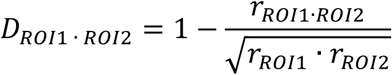

The final matrix containing dissimilarity indices was then used to compute an MDS solution as described previously.

#### Differences between core and extended system representational geometries

In order to quantify differences in representational geometries between areas of the core and extended systems, we divided the pairwise distances between ROIs in the upper triangular RDM into within-system and between-system cells, and converted them back to correlations (by subtracting them from 1). Then, we ran a Linear Mixed-Effect Model on the correlations using *lme4*^54^, fitting a linear model of the form

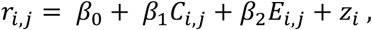

where *i* = 1 … *N* indicates either the subjects for task data (*N* = 33) or the pairwise subjects for hyperaligned movie data (*N* = 55); *j* = 1 … 465 indicates the index of the pairwise correlations between ROIs, *C*_*i,j*_ and *E*_*i,j*_ indicate whether *r*_*i,j*_ is a within-system correlation for the core or extended system respectively, *β*_0_, *β*_1_, *β*_2_ are fixed-effects parameters, and *z*_*i*_ are the subject-level random effects. Using this model, *β*_1_ corresponds to the contrast “Within Core > Between”, and *β*_2_ to the contrast “Within Extended > Between”. After fitting, we performed parametric bootstrapping to obtain 95% bootstrapped confidence intervals on the model parameters.

## Visualization

Volumetric results were visualized using Nilearn ^82^, and projected on template surfaces using AFNI and SUMA ^83,84^.

## Data and code availability

Non-thresholded statistical maps can be found on <u>neurovault.org</u> ^85^ at the following URL: http://neurovault.org/collections/NEUNABLT. All data can be found at http://datasets.datalad.org/?dir=/labs/gobbini/famface/data. The code used for the analyses is available at the following github repository: https://www.github.com/mvdoc/famface.

## Acknowledgments

The authors would like to thank Jim Haxby, Brad Duchaine, the members of the GobbiniLab and HaxbyLab for helpful comments and discussions on this work, and Courtney Rogers for invaluable help with subject recruitment.

## Author contribution

MIG designed the experiment. JDG, JSG, and YOH conducted the experiment. MVdOC, YOH, and JSG analyzed the data. JSG provided analytic tools and critical input to the manuscript. MVdOC, YOH, and MIG wrote the manuscript.

## Footnotes

1 We thank Satrajit Ghosh and Anne Park for sharing the original pipeline.

